# Molecular basis for thermoresponsive protein condensation in plants

**DOI:** 10.64898/2025.12.18.695255

**Authors:** Sunita Pathak, Ananya Chakravarti, Carissa Bersche, Jerelle A. Joseph, Lucia C. Strader

**Affiliations:** The Salk Institute for Biological Studies, La Jolla, CA, USA; Department of Biology, Duke University, Durham, NC, USA; Department of Chemical and Biological Engineering, Princeton University, Princeton, NJ, USA; Omenn-Darling Bioengineering Institute, Princeton University, Princeton, NJ, USA

## Abstract

Plants continuously experience temperature fluctuations that shape their physiology, development, and survival. Yet, how plants sense and adapt to elevated temperatures at the molecular level remains only partially understood. Here, we uncover a sequence-encoded mechanism by which plant proteins directly sense temperature through phase separation. Focusing on prion-like low-complexity domains (PLCDs) in *Arabidopsis thaliana*, we combine cellular imaging, quantitative biophysics, and coarse-grained molecular dynamics simulations to show that numerous PLCDs undergo reversible lower critical solution temperature (LCST)-type transitions. We find that PLCDs that are sufficient and necessary for phase separation confer thermoresponsive variability to full-length proteins. Sequence analysis and simulations reveal that thermoresponsiveness is encoded in compositional heuristics, with sufficient PLCDs enriched in aliphatic and aromatic residues, enabling accurate prediction and rational tuning of condensation thresholds. Engineered variants with altered sequence composition exhibit predictable shifts in thermoresponsive behavior, confirming a direct, sequence-level encoding of temperature sensitivity. Our findings establish LCST-driven phase separation as a fundamental mechanism of thermoresponsiveness in plants and provide a molecular framework for designing synthetic biomolecules with programmable temperature responses.

---

Plants are complex biological systems with a remarkable capacity to sense and adapt to fluctuating daily and seasonal temperatures. As ectothermic organisms, their cellular temperature closely tracks the surrounding environment, making nearly all aspects of plant physiology—from growth and development to photosynthesis, flowering, and reproduction—highly sensitive to thermal conditions. Indeed, temperature is a critical cue sensed by plants to drive these specific biological processes [1]. Each plant species therefore operates within an optimal temperature range that shapes its geographic distribution, ecological diversity, and overall adaptability. Yet, plants frequently encounter temperatures outside this range, with elevated heat posing a particularly critical challenge. Heat stress disrupts normal growth of plants, compromising both plant quality and productivity [2, 3]. In the context of ongoing climate change and projected global temperature increases, understanding the molecular mechanisms that govern plant responses to heat stress is urgent for safeguarding crop yield, ensuring food security, and preserving ecosystem biodiversity. Moreover, emerging synthetic biology approaches offer powerful strategies to engineer resilience and mitigate the detrimental effects of rising temperatures [4].

Plants perceive and respond to temperature fluctuations through a diverse set of cellular and molecular mechanisms. Elevated temperatures result in thermomorphogenesis, a growth response which includes hypocotyl elongation, leaf hyponasty, changes in leaf morphology, altered flowering time, and root elongation [1]. Stressful elevated temperatures can also trigger mechanisms of both basal and acquired thermotolerance, which result in physiological changes that enable persistence under elevated temperatures [5, 6]. These adaptive strategies are mediated in part by thermosensory pathways, such as the well-characterized phytochrome B, PIF7 and ELF3 pathways [7–9] and the activation of heat shock transcription factors that regulate protective chaperone networks [10, 11]. Beyond these well-studied pathways, emerging evidence highlights a new layer of thermal sensing at the molecular level: the phase separation of proteins in response to temperature [9, 12–14]. This temperature-dependent condensation represents a new frontier in understanding how plants reorganize their molecular machinery to cope with elevated temperatures.

Importantly, in certain organisms—particularly within the plant and fungal kingdoms—protein condensation has emerged as a central mechanism for sensing environmental stressors [15]. In response to temperature changes, phase separation of proteins can drive the formation of biomolecular condensates, which are membraneless organelles sustained by dynamic interactions between proteins and/or nucleic acids. The ability of these condensates to assemble and disassemble in a multimeric fashion enables cells to rapidly reorganize their molecular machinery in tune with environmental cues. In plants, temperature-regulated condensation tunes thermomorphogenesis and thermotolerance [9, 12–14, 16, 17]. Further, heat-induced condensation gives rise to cytoplasmic stress granules, processing bodies [18, 19] and chloroplastic condensates [20], all of which play pivotal roles in maintaining cellular homeostasis under stress. Although significant progress has been made in characterizing these phenomena, a deep mechanistic understanding of how temperature triggers and regulates protein condensation in plants remains elusive. Unraveling these molecular principles is essential not only for advancing our fundamental knowledge of cellular stress responses but also for developing strategies to enhance plant resilience in a warming climate.

Several condensate-mediated mechanisms have been proposed by which heat triggers adaptive stress responses in cells. A prevailing view has been that heat-induced denaturation or damage generates misfolded proteins, which in turn drive condensate formation [21]. Such mechanisms have been described for ribosomal subunits, RNA-binding proteins, and other stress granule-associated proteins [21]. Central to this model is the role of molecular chaperones, which sequester and refold misfolded proteins to restore proteostasis [22] alongside phosphorylation-dependent signaling events that initiate downstream stress responses [23]. However, emerging evidence points to an alternative paradigm: rather than forming condensates as a consequence of damage, certain proteins can directly sense temperature shifts and undergo phase separation [9, 14]. This raises the possibility that temperature response is, at least in part, encoded within the intrinsic sequence features of proteins recruited to condensates. In this context, a critical question emerges—can sequence-encoded, temperature-responsive transitions in proteins govern heat-induced condensation?

Across species, many sequences that drive condensation are enriched in prion-like low complexity domains (PLCDs), which contain amino acid regions characterized by intrinsic disorder and low sequence complexity [24–26]. PLCDs can encode distinct modes of phase behavior. In mammals, numerous PLCD-containing proteins undergo upper critical solution temperature (UCST)-type transitions [27], where condensates form at lower temperatures and dissolve upon heating. In contrast, in ectothermic organisms such as plants and fungi, PLCDs in proteins like ELF3 [9] and Pab1 [28] have been shown to mediate lower critical solution temperature (LCST)-type transitions, in which condensates assemble upon heating. Notably, plants such as *Arabidopsis thaliana* encode a large repertoire of proteins with PLCDs [29, 30] raising the question of whether these domains act broadly as sequence-encoded sensors of temperature to modulate plant growth and physiology.

Here, we combined experimental and computational approaches to investigate whether PLCD-containing Arabidopsis proteins encode thermoresponsive behavior and to identify the sequence features that drive it. We first used protoplast-based experimental assays to test the response of PLCDs at varying temperatures and found that many are sufficient to mediate condensation; however, these displayed notable variation in thermoresponsive behavior. Strikingly, numerous PLCDs encoded LCST-type transitions, and their sufficiency and necessity for phase separation linked directly to temperature sensitivity. We found that PLCD sufficiency to drive phase separation was directly encoded in the variability of condensation response: PLCDs that were sufficient for phase separation conferred high thermoresponsive variability to the full-length protein. Residue-level coarse-grained molecular dynamics simulations further revealed that PLCD thermoresponsiveness was determined by specific sequence features, with sufficient PLCDs having a higher fraction of aliphatic and aromatic residues. Leveraging these heuristics, we predicted PLCD behavior, engineered mutants with altered thermoresponsiveness, and validated these predictions experimentally. Together, our findings establish a direct connection between LCST-type transitions and plant temperature responses and lay a foundation for harnessing sequence-encoded condensation principles in synthetic biology.

## Results

### PLCD-containing proteins in Arabidopsis exhibit variability in heat-induced condensation behavior

To investigate how PLCDs contribute to thermoresponsive condensation, we selected *Arabidopsis thaliana* as our model system given its well-annotated genome and its extensive use in temperature response studies. Using the Arabidopsis Columbia-0 (Col-0) reference genome, we scanned 27,745 proteins with PLAAC (Prion-like Amino Acid Composition; setting minimum PLCD length = 60, *S. cerevisiae* background frequencies, α = 100) [31]. This analysis identified 482 proteins containing predicted PLCDs (∼1.7% of proteins; Supplemental Table 1), consistent with prior studies [29]. Gene ontology enrichment revealed that these PLCD-containing proteins are enriched in RNA and DNA binding, protein dimerization, and transcriptional regulation (Fig. 1a); these functional categories have previously been associated with phase separation in yeast and human PLCDs [32–35]. Based on these observations, we hypothesized that Arabidopsis PLCD-containing proteins may also undergo phase separation in response to temperature changes.

**Figure 1.**
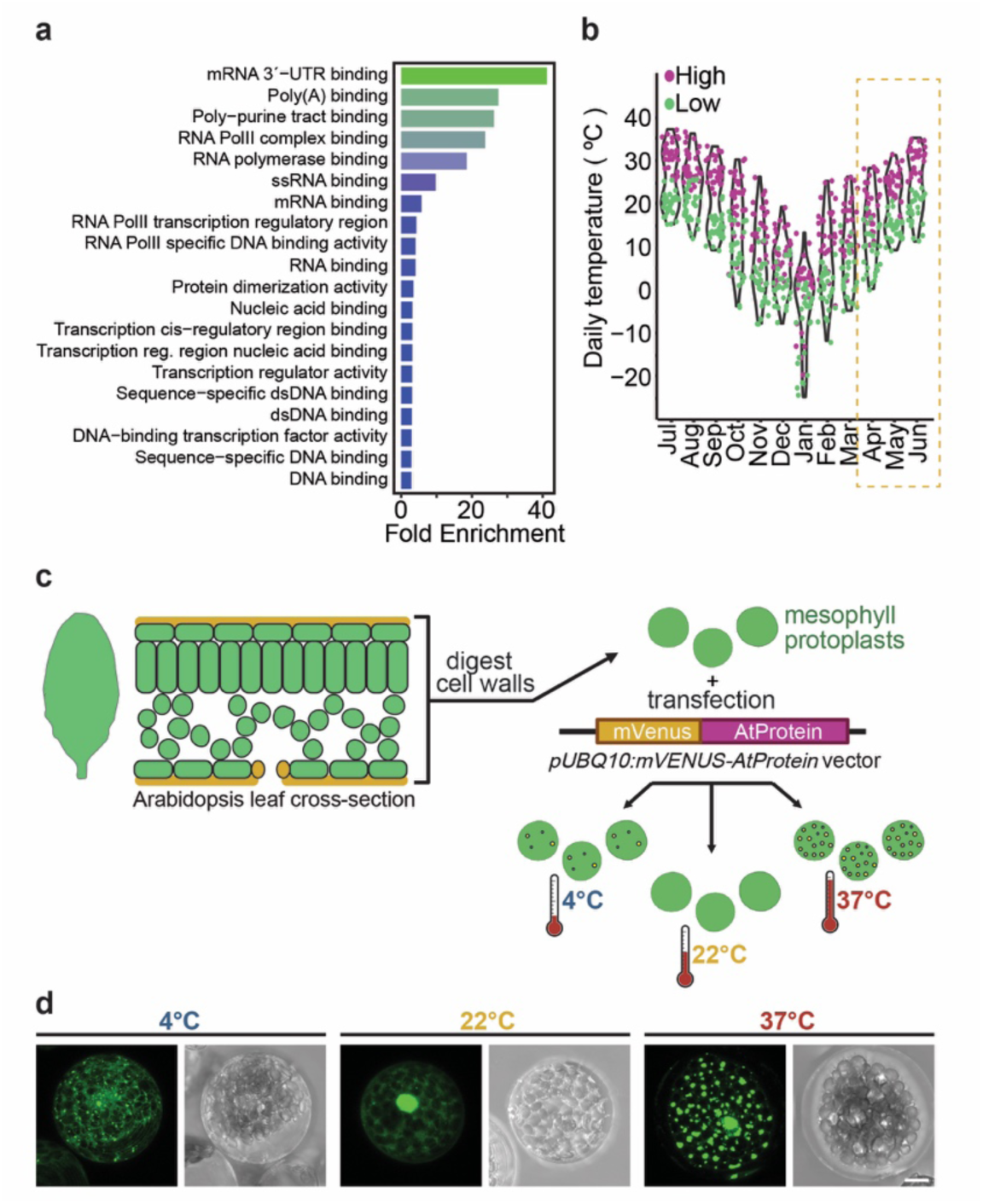
A high-throughput protoplast-based assay enables examination of thermoresponsive condensate formation. (a) Bar graph showing the top 20 enriched GO category (molecular function) of predicted PLCDs with fold enrichment shown on x-axis and GO categories on y-axis. Plot was generated using ShinyGO (https://bioinformatics.sdstate.edu/go/). (b) Violin plot representing the daily temperature fluctuation in Columbia, Missouri through the year 2024. Orange color (high temperature) and cyan color (low temperature). The highlighted rectangular area represents the optimum growing season of *Arabidopsis thaliana*. Data collected from https://weather.com. (c) Arabidopsis mesophyll cells were isolated by enzymatic cell wall digestion, allowing transfection with protein expression plasmids. Protoplasts were then incubated under cold (4 °C), normal (22 °C), or heat (37 °C) conditions for 30 minutes. (d) The system recapitulates known stress granule dynamics, as demonstrated by the condensation behavior of the stress granule protein RBP45B. The images show fluorescent signal (YFP) and the corresponding bright field images taken at 4 °C (left), 22 °C (middle) and 37 °C (right). Scale bar = 10 µm.

Because Arabidopsis completes its growth cycle during spring months when temperatures range from ∼4–37 °C (Fig. 1b), we sought to probe condensate formation across this physiological range. Although laboratory growth is typically performed at 22 °C, we included 4 °C (low temperature), 22 °C (baseline), and 37 °C (heat stress) to examine thermoresponsive behavior. We employed a protoplast-based assay (Fig. 1c,d), in which leaf mesophyll cells were enzymatically digested to generate protoplasts amenable to transient DNA uptake. Candidate PLCD-containing proteins were expressed as N-terminal mVenus fusions under the *pUBQ10* promoter, enabling rapid and high-throughput evaluation of condensate formation. Protoplasts were incubated for 16 h at 22 °C, followed by a 30-minute shift to 4, 22, or 37 °C prior to imaging.

From the pool of 482 proteins (Supplemental Table 1), we tested 54 PLCD-containing candidates selected based on cDNA availability from the Arabidopsis Biological Resource Center (Supplemental Table 2). These candidates spanned across the list of molecular functions for predicted PLCD-containing proteins (Supplemental Table 2). We quantified condensate formation propensity by measuring the percentage of cells displaying fluorescent puncta (out of all cells expressing the protein) at each temperature. Whereas mammalian PLCD-containing proteins typically undergo upper critical solution temperature (UCST)-type transitions (condensation at lower temperatures)[36], we observed that most Arabidopsis PLCD-containing proteins exhibited lower critical solution temperature (LCST)-like behavior, condensing at higher temperatures (Fig. 2).

**Figure 2.**
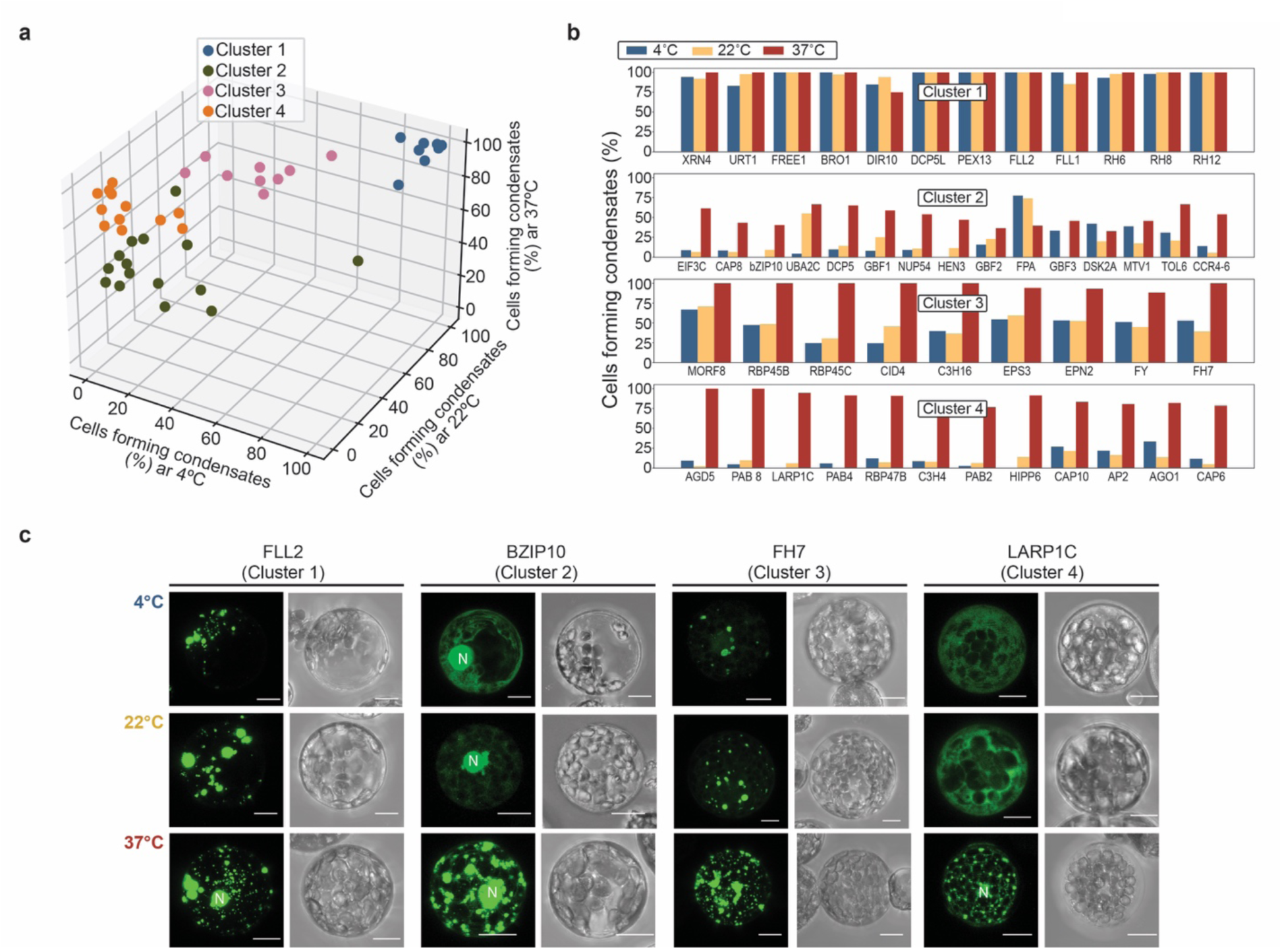
PLCD-containing proteins exhibit variability in thermoresponsive behavior. (a) *k*-means clustering with silhouette analysis identifies four distinct clusters of condensation behavior across 4 °C, 22 °C, and 37 °C. Proteins were assigned to the nearest cluster centroid based on Euclidean distance. Cluster 1 proteins display high condensation at all temperatures. Cluster 2 proteins show low condensation at 4 °C and 22 °C and moderate condensation at 37°C. Cluster 3 proteins exhibit moderate condensation at 4 °C and 22 °C and high condensation at 37 °C. Cluster 4 proteins show low condensation at 4 °C and 22 °C and high condensation at 37°C. (b) Percentage of cells displaying mVenus-AtProtein puncta at each tested temperature. For each temperature, in average 50 cells were examined at each temperature. The protoplast count from two replicates were combined to plot the graph. (c) Representative protoplast images from each cluster at 4 °C, 22 °C, and 37 °C. FLL2 from Cluster 1, bZIP10 from Cluster 2, FH7 from Cluster 3 and LARP1C from Cluster 4. The images show the YFP signal (Left) and bright filed image (right) of each protein. Scale bar = 10 µm. N indicates the nucleus.

Furthermore, we observed that most of these full-length PLCD-containing proteins (48 out of 54) exhibited temperature responsive condensation behavior, which showed a variable pattern of condensation. To systematically classify these patterns of condensation, we applied *k*-means clustering, using silhouette scores to determine the optimal cluster number (Fig. 2a). Four distinct clusters emerged: Cluster 1, high condensation across all tested temperatures; Cluster 2, minimal condensation at 4 and 22 °C with moderate condensation at 37 °C; Cluster 3, moderate condensation at 4 and 22 °C with high condensation at 37 °C; and Cluster 4, minimal condensation at 4 and 22 °C with high condensation at 37 °C (Fig. 2b).

As 37 °C represents a physiologically relevant heat stress, Clusters 2–4 contain many proteins that form condensates specifically in response to elevated temperature, such as DCP5 [37], MORF8 [20], and RBP47B [38], consistent with LCST-type transitions. Although, Cluster 1 proteins form condensate across all tested temperatures, they still display temperature responsive change in condensate morphology (SI Fig. S1).This behavior resembles the dynamics reported for processing bodies[39] and accordingly, cluster 1 includes several known processing body localized proteins like RH6, RH8 and RH12[40].

Overall, these results reveal striking variation in the thermoresponsive behaviors of Arabidopsis PLCD-containing proteins. They also raise two central questions: 1) what sequence features drive this variability in phase separation, and 2) to what extent is variability in phase separation encoded by the PLCDs themselves?

### Those PLCDs sufficient for condensation underlie thermoresponsive variability in full-length proteins

To test whether PLCDs directly drive thermoresponsive condensation, we selected 20 representative proteins from our dataset and examined both isolated PLCD regions and full-length variants with PLCDs deleted. This design allowed us to assess whether PLCDs are sufficient to form condensates and/or are necessary for condensation in the context of the full-length protein.

From this analysis, three distinct categories emerged. Category A proteins formed condensates upon heat stress regardless of PLCD deletion (Fig. 3a), whereas their isolated PLCDs failed to condense, suggesting that the PLCDs in this group were neither sufficient nor required for condensation. In Category B, isolated PLCDs robustly condensed at elevated temperature, and removal of these domains abolished or severely diminished condensation in the full-length protein (Fig. 3b), demonstrating that the PLCDs were both sufficient and necessary. Finally, in Category C, isolated PLCDs alone failed to condense, but deletion of these domains in the full-length protein strongly reduced condensate formation (Fig. 3c). Thus, while PLCDs in this group were not sufficient, they were nonetheless necessary for thermoresponsive condensation.

**Figure 3.**
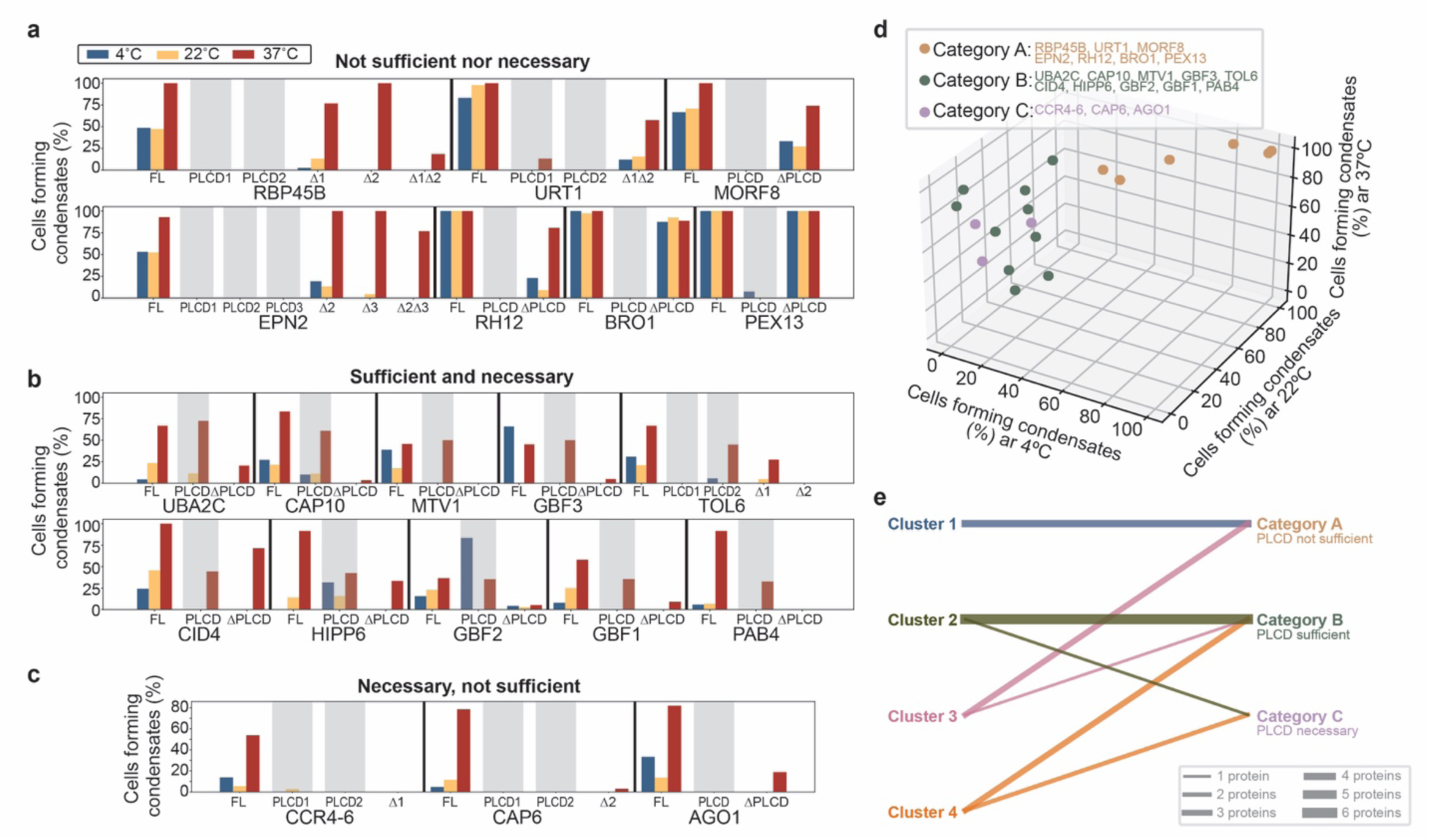
PLCDs of proteins display varying degrees of sufficiency in thermoresponsive condensation, which correlates to the thermoresponsive variability of full-length proteins. (a,b,c) Bar plots representing the percentage of cells displaying mVenus-AtProtein puncta, mVenus-AtProtein^PLCD^ (just PLCD) puncta and mVenus-AtProtein^ΔPLCD^ (deleted PLCD) puncta at 4 °C, 22 °C, and 37 °C in three distinct categories of PLCD sufficiency: (a) Category A PLCDs are neither sufficient nor necessary: full-length proteins condense upon heat stress even when PLCDs are deleted, while isolated PLCDs fail to condense. (b) Category B PLCDs are both sufficient and necessary: isolated PLCDs robustly condense at elevated temperature, and deletion of these domains diminishes condensation in the full-length protein. (c) Category C PLCDs are necessary but not sufficient: isolated PLCDs alone do not condense, but their deletion strongly reduces condensation in the full-length protein. (d) *k*-means clustering with silhouette analysis on the full-length proteins shows that Categories B and C have higher thermoresponsive variability than Category A, indicating that PLCD sufficiency maps to higher variability in the full-length protein. Higher and lower variability was defined by *k*-means clustering on percentage of cells with condensates across 4°C, 22°C, and 37°C. Proteins were assigned to the nearest cluster centroid based on Euclidean distance. (e) All Category A proteins map to Clusters 1 and 3 (low variability), whereas nearly all Category B and C proteins map to Clusters 2 and 4 (high variability). Line thickness corresponds to number of proteins, as indicated in the legend.

Having established these functional categories, we next asked whether PLCD sufficiency and necessity could explain the variation in thermoresponsive behaviors observed in full-length proteins. Using *k*-means clustering with silhouette analysis, we classified condensation profiles of the full-length proteins (Fig. 3d). Proteins in Categories B and C were characterized by a marked increase in condensation as temperature rose—i.e., high variability in thermoresponsive behavior. By contrast, Category A proteins were characterized by stable condensation across temperatures. These results reveal a key principle: when PLCDs are sufficient or necessary, the corresponding full-length proteins display high thermoresponsive variability, whereas proteins in which PLCDs are dispensable show little change across temperatures.

Finally, we compared the PLCD-based functional categories with the global clustering of full-length proteins. Remarkably, all Category A proteins mapped to Clusters 1 and 3 (low variability), whereas nearly all Category B and C proteins mapped to Clusters 2 and 4 (high variability) (Fig. 3e). Thus, the condensation behavior of isolated PLCDs is predictive of thermoresponsive variability in the full-length protein.

Together, these findings establish a direct and previously unrecognized link between PLCD sufficiency/necessity and the degree of thermoresponsive phase behavior. This highlights PLCDs as drivers of heat-induced condensation, providing a mechanistic explanation for the diversity of phase separation behaviors in Arabidopsis proteins. Following this, the critical question became: what features determine whether a PLCD is sufficient to drive thermoresponsive phase separation?

### PLCD sufficiency and necessity in heat-induced phase separation is encoded in sequence features

To uncover the molecular drivers of PLCD-mediated thermoresponsive condensation, we turned to molecular simulations and computational modeling to gain quantitative insight. While atomistic simulations provide high resolution, they are prohibitively expensive for studying phase separation, which requires modeling large ensembles of interacting proteins over extended length and time scales [41]. Residue-level coarse-grained approaches strike a balance between resolution and efficiency, enabling the study of protein dynamics and condensate formation under physiologically relevant conditions. In particular, the Mpipi-T model has been validated as a quantitatively accurate residue-level coarse-grained framework for predicting both single-molecule behavior and collective phase separation as a function of temperature, especially for heat-induced LCST-type transitions [42]. Notably, Mpipi-T has been shown to recapitulate the LCST phase behavior of ELF3, a PLCD-containing protein in the Arabidopsis evening complex that is central to thermosensory regulation [9]. This result provided strong confidence that the model would be well-suited for probing PLCD-driven condensation in our system.

Using Mpipi-T [42], we first simulated individual PLCDs and computed the Flory scaling exponent from the radius of gyration (Fig. 4a). From these simulations, we identified the coil-to-globule transition temperatures, which quantify the temperature at which PLCDs undergo intrachain collapse. We found that PLCDs sufficient for thermoresponsive phase separation transitioned to more compact globule states at lower temperatures than those that were insufficient, consistent with enhanced LCST-type behavior (Fig. 4c, SI Fig. S2). Extending this analysis to multi-chain simulations using the direct coexistence approach (Fig. 4b), we determined the dense-phase densities across temperatures. Sufficient PLCDs achieved higher dense-phase packing as temperature increased compared to insufficient PLCDs (Fig. 4d, SI Figs. S3–S5). Together, these results reveal that sufficiency reflects a greater propensity for chain compaction at elevated temperatures and enhanced assembly into condensates.

**Figure 4.**
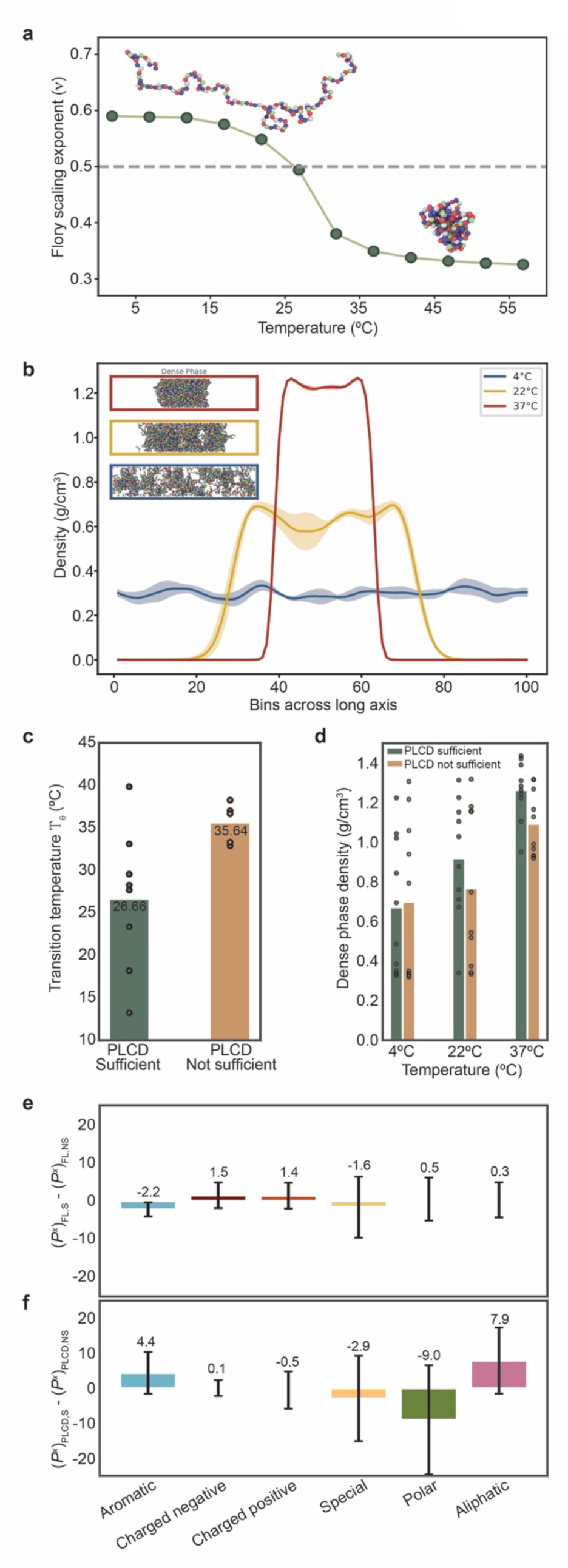
Sequence drivers govern PLCD thermoresponsive condensation. Molecular dynamics simulations are performed using the Mpipi-T model to simulate single chain and multiple chain protein systems. (a) For single chain systems, the Flory scaling exponent (ρ) is calculated from the simulated protein’s radius of gyration to determine the transition from coil (expanded) to globule (compact) state. The dashed gray line indicates ρ = 0.5, which is the coil-to-globule transition. (b) For multiple chain systems, direct coexistence simulations are performed to obtain the density profile along the long axis of the simulation box at various temperatures, from which the dense-phase density is extracted. The results from the simulations show that PLCDs sufficient for condensation display (c) lower coil-to-globule transition temperatures in single-chain simulations and (d) higher dense phase densities under heat stress in multi-chain simulations. Bars represent the mean across proteins, and individual proteins are shown as points. These behaviors reflect underlying sequence heuristics: (e) full-length proteins do not differ significantly in residue composition based on PLCD sufficiency, but (f) sufficient PLCDs are enriched in aliphatic and aromatic residues and depleted in polar residues compared to insufficient PLCDs. Bars represent the mean residue fraction across sequences, and error bars represent the standard deviation.

Because Mpipi-T is parameterized with residue-level chemical specificity [42], we hypothesized that PLCD sufficiency could be explained by sequence composition. PLCDs in mammalian systems are enriched in aromatic and polar residues, thus we wanted to know what distinct sequence features govern thermoresponsive condensation in plants. When we compared residue fractions across full-length Arabidopsis proteins, we did not find any significant differences in residue composition (Fig. 4e). However, when we focused specifically on PLCD sequences, a pattern emerged: sufficient PLCDs were enriched in aliphatic and aromatic residues and depleted in polar residues compared to insufficient PLCDs (Fig. 4f).

These findings suggest that PLCD sufficiency for heat-induced condensation is encoded at the sequence level and is linked to distinct residue biases. Importantly, these sequence-level features of PLCDs translate into variability in the thermoresponsive behavior of the full-length protein.

This work therefore uncovers a set of predictive heuristics that connect amino acid composition to phase separation outcomes, advancing our mechanistic understanding of thermosensing in Arabidopsis and offering principles that may extend across other plant species.

### Sequence heuristics can be leveraged to design mutants with tunable thermoresponsive phase separation

Having identified sequence heuristics that govern PLCD sufficiency for thermoresponsive condensation, we next asked whether these rules could be leveraged predictively to engineer mutants with altered phase behavior. Specifically, we focused on aliphatic residues as putative drivers of Arabidopsis PLCD LCST-type phase separation, which is driven primarily by entropic effects [42–44]. Using Mpipi-T, we designed variants for the proteins with the lowest and highest aliphatic residue fractions (Supplemental Table 3). For each protein, we generated three classes of mutants: Variant Set 1, in which polar or aromatic residues were mutated to aliphatic residues (A, V, M; 50% aliphatic enrichment); Variant Set 2, in which only polar residues were mutated to aliphatic residues (A, V, M; 50% aliphatic enrichment); and Variant Set 3, in which polar residues were mutated to yield 25% aliphatic (A, V, M) and 25% aromatic (Y) enrichment, totaling 50% combined (Fig. 5a, for example). The specific residues were chosen because A, V, M, and Y were the most frequently represented aliphatic and aromatic residues in the native PLCDs we analyzed. The 50% enrichment target was inspired by aliphatic enrichment of elastin-like polypeptides designed to exhibit robust LCST behavior [44]. Based on our heuristics, we hypothesized Variant Set 3 would show the strongest heat-induced condensation, followed by Variant Set 2 and then Variant Set 1.

**Figure 5.**
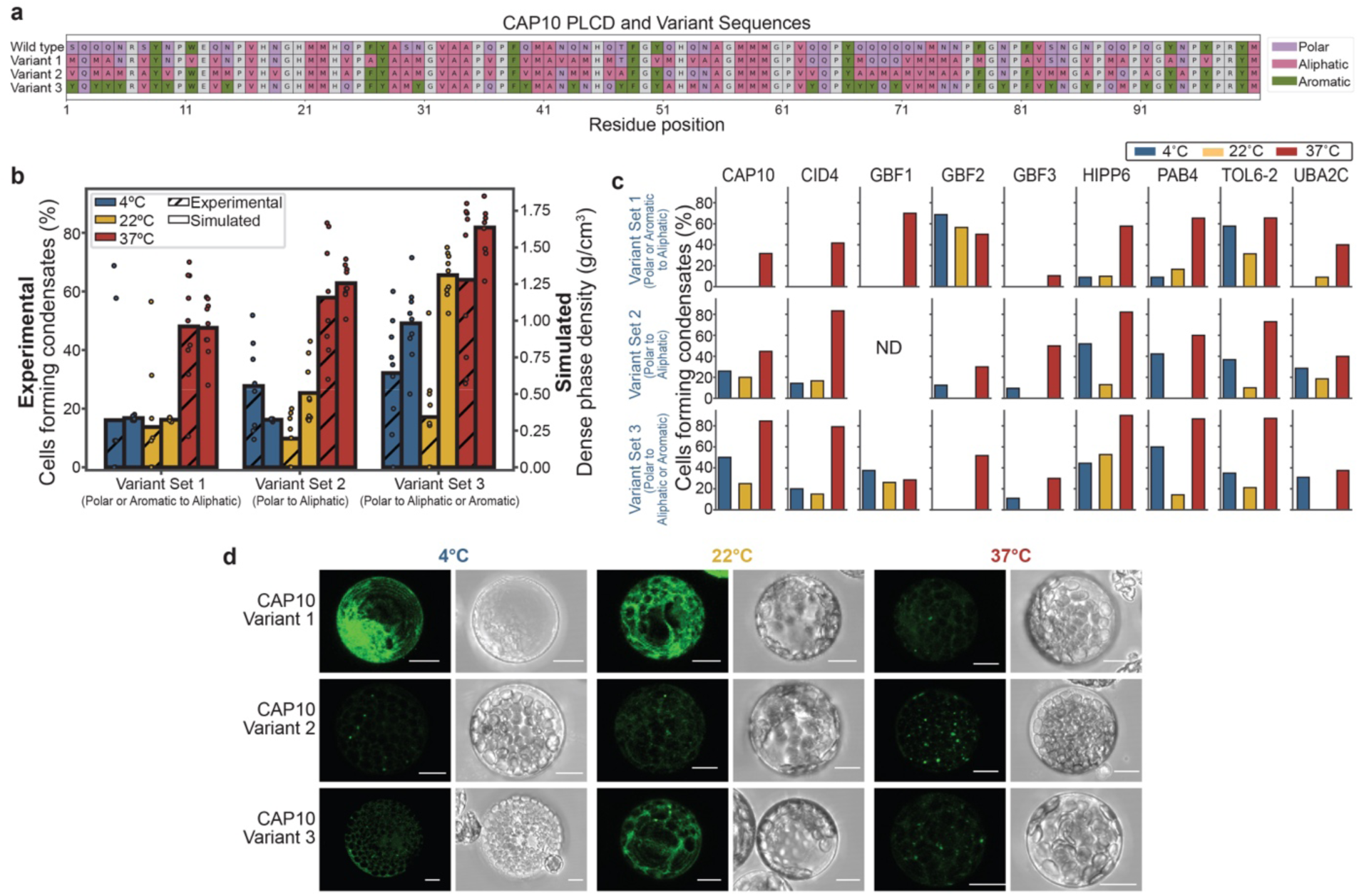
Sequence heuristics enable engineering of protein variants with tunable thermoresponsive condensation. Nine protein variants were designed in three sets: Set 1—polar or aromatic residues mutated to achieve 50% aliphatic enrichment; Set 2—polar residues mutated to achieve 50% aliphatic enrichment; and Set 3—polar residues mutated to achieve 25% aliphatic and 25% aromatic enrichment. (a) The wild-type CAP10 sequence and its three designed variants are shown as representative examples. (b) Direct coexistence simulations using the Mpipi-T model were used to determine dense phase densities of variants at 4 °C, 22 °C, and 37 °C. The simulations predicted that Set 3 variants, enriched in both aliphatic and aromatic residues, would exhibit the strongest heat-induced condensation. Experimental high-throughput protoplast-based assays were performed at 4 °C, 22 °C, and 37 °C for each of these computationally designed variants to quantify the percentage of cells with condensates. The experimental results recapitulated the simulation predictions (Pearson r = 0.726, p = 0.02686), validating that polar-to-aliphatic and aromatic substitutions maximize condensation propensity under heat stress. Bars represent the mean across variants in a set, and individual variants are shown as points. (c) Experimental quantification of the percentage of cells forming condensates at 4 °C, 22 °C, and 37 °C for all designed variants. An average of 30 cells were imaged and quantified. (d) Representative microscopy images show condensate formation for CAP10 protein variants under the three temperature conditions. YFP signal image (Left) and bright field image (right) for each temperature and variants. Scale bar = 10 µm.

Mpipi-T simulations with confirmed this prediction (Fig. 5b, SI Figs. S6–S8). Variant Set 1 showed minimal condensation at 4 °C and 22 °C but increased condensation at 37 °C. Variant Set 2 exhibited low condensation at 4 °C, moderate condensation at 22 °C, and high condensation at 37 °C. Variant Set 3 displayed the highest condensation propensity overall, with substantial condensation already at 22 °C and maximal condensation under heat stress (37 °C). These results directly align with the residue-level biases we identified: enrichment of aliphatic and aromatic residues, coupled with depletion of polar residues, enhances thermoresponsive phase separation of PLCDs.

To validate these predictions, we experimentally tested all variants in our protoplast-based assay. The experimental results closely mirrored the simulations: Variant Set 3 showed the strongest heat-induced condensation, followed by Variant Set 2 and then Variant Set 1 (Fig. 5b and c). This convergence between simulation and experiment underscores the predictive power of our sequence heuristics.

Together, these findings demonstrate that PLCD phase behavior can be rationally tuned by manipulating amino acid composition, providing direct evidence that sequence-level rules are sufficient to guide the design of thermoresponsive condensates in plants. The strong agreement between computational predictions and experimental validation highlights the robustness of our framework and opens the door to engineering synthetic proteins with tailored phase behaviors for biotechnology and synthetic biology applications.

## Discussion

In this work, we uncover fundamental principles underlying thermoresponsive condensation in plants. Using a high-throughput *Arabidopsis thaliana* protoplast-based assay, we systematically profiled the phase separation behavior of PLCD-containing proteins across a range of temperatures. We observed striking variability in condensation behavior, with most proteins exhibiting heat-induced condensation consistent with LCST-type transitions. This variability mapped directly to the sufficiency or necessity of PLCDs: proteins in which PLCDs were sufficient or necessary for condensation displayed markedly greater thermoresponsive variability in the full-length protein. Leveraging residue-resolution coarse-grained molecular dynamics simulations, we identified sequence-encoded heuristics as the molecular drivers of this behavior. Specifically, PLCDs sufficient for heat-induced condensation were enriched in aliphatic and aromatic residues and depleted in polar residues, linking amino acid composition to condensate propensity. These sequence heuristics not only explained natural variation but also enabled design: by engineering mutants with targeted amino acid fractions, we were able to directly manipulate the thermoresponsive condensation behavior of proteins. Remarkably, the trends predicted by simulations were recapitulated experimentally, validating that variants in which polar residues were mutated to aliphatic and aromatic residues had the highest propensity for condensation under heat stress. This convergence between computational prediction and experimental validation underscores the robustness of our framework and provides a foundation for considering the broader implications of these findings.

Together, these results reveal a direct sequence-to-phenotype connection for plant PLCDs and highlight their central role as drivers of heat-induced protein condensation. This finding allows us to advance a previously overlooked hypothesis: that an intrinsic component of the plant temperature response is grounded in LCST-type phase behavior. Extending beyond plants, these principles may illuminate the molecular logic of thermosensing across ectothermic organisms—an especially timely question in the context of accelerating climate change.

Beyond basic cell biology, our work establishes a blueprint for engineering thermoresponsive proteins. The sequence heuristics and phenotype mappings we uncovered provide a rational framework for designing synthetic biopolymers with tailored heat-induced condensation properties. Looking forward, the computational–experimental pipeline we developed can be extended to generate novel variants for use as biosensors, smart materials, environmental remediators, and agricultural tools, underscoring both the conceptual and translational potential of this work.

## Limitations

While establishing the promising avenues of our findings, it is also important to acknowledge the limitations. This study focused on a subset of PLCD-containing proteins, and expanding the analysis to a broader set will be important for deriving more generalizable principles that connect sequence features to condensation propensity. On the computational side, we assessed condensate density as a proxy for phase separation propensity and compared this with the percentage of cells exhibiting condensates in experiments. Although informative, this comparison could be strengthened by incorporating additional morphological descriptors, such as condensate size, number, and total area, which collectively may provide a more nuanced link between simulation and experiment. Establishing a robust framework to directly map computational dense-phase density to experimentally measured condensate morphology will require not only methodological development but also more controlled experimental systems, as our current protoplast assay is subject to confounding factors such as other biomolecules and small environmental fluctuations. Our study does not explore the link between our engineered variants and biological function, which will require long-term assessment. Finally, our study was limited to a single Arabidopsis ecotype (Col-0). It will be valuable to investigate other ecotypes and protein homologs, as natural variation in thermoresponsive behavior may reflect geographic adaptation and provide further insights into how PLCD-driven condensation contributes to environmental fitness.

## Materials and Methods

### Plant growth

Arabidopsis seeds (Col-0) were surface sterilized with a bleach solution (10% Clorox + 0.01% Triton X-100) for 10 minutes and washed in sterile water three times prior to resuspension in 0.1% agar. The suspended seeds were then stratified at 4 ℃ for 2 days. Stratified seeds were then plated on plant nutrient (PN) medium solidified with 0.6% agar. The plated seeds were grown for 10 days at 22 °C under continuous light which were transplanted in soil and grown for an additional week under continuous light at 22 °C.

### Protoplast isolation and transfection

Protoplasts were isolated from 17-day-old Arabidopsis Col-0 plants using the tape method as described in [45] with slight modifications. The peeled leaves were incubated for 2 hours in the enzyme solution with continuous shaking at 100x rpm. Protoplast transfection was performed as described in [45] with slight modifications. 10 µg plasmid DNA was added to 100,000 protoplasts suspended in 200 µL MMG solution. 210 µL volume of PEG solution was then added to the protoplasts in MMG and incubated at room temperature for 15 min. The protoplasts in the PEG solution were then washed twice with 800 µL W5 solution and resuspended in 500 µL W5 solution. Transfected protoplasts were incubated at 22 °C for 12-16 hours to allow for protein expression. Subsequently, transfected protoplasts were incubated at 4 °C, 22 °C, or 37 °C for 30 minutes prior to data acquisition.

### Vector construction

Full-length CDS was obtained from Arabidopsis Biological Resource Center and amplified using Platinum SuperFi II DNA polymerase (Invitrogen) and cloned into pENTR/D-TOPO (Invitrogen) to create pENTR-D-full-length CDS using the primers listed in Supplemental Table 4. Predicted PLCD regions were amplified from the CDS and cloned into pENTR/D-TOPO (Invitrogen) using the primers listed in Supplemental Table 4 to generate pENTR-d-PLCD constructs. To generate pENTR-d-noPLCD constructs, PCR was performed on the pENTR-D-full length CDS clones using the primers listed in Supplemental Table 4. The obtained PCR product was then gel purified and treated with DpnI. These PCR products, containing 15-bp overlap regions, were directly transformed into NEB5α cells for homologous recombination [46]. The full-length, PLCD and no PLCD entry vectors were then recombined into pLCS101 (pUBQ10: mVenus-GW) vector [47] to generate *pUBQ10:mVenus-full-length*, *pUBQ10:mVenus-PLCD* and *pUBQ10: mVenus-no PLCD* plasmids. These plasmids were isolated using ZymoPURE II plasmid midiprep kit (Zymo Research) to be used in the aforementioned protoplast transfection protocol.

### Microscopy

Protoplasts were mounted on a glass slide with a silicon isolator (Electron Microscopy Science, Cat. # 70337-20) between the slide and cover slip to confine the protoplasts without compression. Vacuum grease was used to attach the isolator to the slide and cover slip.

Protoplasts were imaged with a Leica SP8 confocal equipped with a HyD detector. Samples were excited with a 512 nm laser and emission collected through a 40x water immersion lens. Z-stacks of each protoplast were acquired, and the maximum projection of images were analyzed.

### *k*-means clustering and silhouette analysis

To classify proteins based on their temperature-dependent condensation behavior, *k*-means clustering was performed. Specifically, values for condensate formation at 4 °C, 22 °C, and 37 °C were standardized using the StandardScaler function in *scikit-learn* [48] to ensure equal weighting across features. The optimal number of clusters was evaluated by computing silhouette scores across *k* values ranging from 2 to 7. The silhouette coefficient, which quantifies the cohesion and separation of clusters, was averaged across all proteins for each *k*. The value of *k* that yielded the highest average silhouette score was selected as the optimal number of clusters. Following this, *k*-means clustering was performed with the optimal cluster number and a random state = 0 to allow reproducibility, and proteins were assigned to their respective groups. Results were visualized using 3D scatterplots generated with the *plotly* package (https://plotly.com/) enabling interactive exploration of cluster assignments.

### Mpipi-T approach

Mpipi-T is a residue-resolution coarse-grained model [42]. In this approach, each amino acid is represented as a unique bead (with defined size, charge, mass). The potential energy between proteins is computed as sum of bonded interactions and non-bonded interactions. To represent bonded interactions, beads are connected via harmonic springs. Non-bonded interactions encompass long-ranged electrostatics, which are modeled via Coulomb term with a temperature-dependent Debye-Hückel screening, and short-ranged pairwise contacts that are also temperature-dependent. The non-bonded interactions in the model were parameterized by combining atomistic free energy of solvation data of residue pairs with experimental cloud point measurements. This parameterization strategy allows Mpipi-T to capture both UCST and LCST-type phase behaviors. Mpipi-T has been validated against experimental data, showing quantitative predictions for both UCST and LCST sequences.

### Dense phase density computations

Dense phase densities were calculated using direct coexistence simulations in the slab geometry via the Mpipi-T residue-resolution coarse-grained model [42]. Each system was simulated in the canonical ensemble with a Langevin thermostat (relaxation time = 5 ps). Simulations consisted of a 50 ns equilibration phase followed by a 400 ns production run with a timestep of 10 fs. The dense and dilute phases were identified at equilibrium, and the dense phase density was quantified from the mass of residues in the high-density region normalized by simulation volume.

### Coil-to-globule transition computations

The coil-to-globule transition was quantified by computing the Flory scaling exponent from the radius of gyration, following procedures described in [42]. Canonical ensemble simulations with a Langevin thermostat (relaxation time = 5 ps) were performed with a 0.5 μs equilibration period followed by 3 μs of production, using a timestep of 10 fs. Simulations were conducted in 10 K increments across a temperature range of 280 to 370 K. The radius of gyration was recorded every 1,000 timesteps (0.01 ns), and the Flory scaling exponent was calculated to identify the coil-to-globule transition temperature.

### Unique biological materials

Unique biological materials are available from the corresponding author upon request.

## Data availability

All data are either reported in this manuscript or are available upon request from the corresponding authors.

## Code availability

Code developed and used in this manuscript are available on the GitHub repository: github.com/josephresearch/plant_lcst.

## Acknowledgements

We thank Ryan Bitter, Billy Cao, Suresh Damodaran, Anita Donlic, Jhonny Figueroa, Libby Gilmore, David Korasick, and Se-Hwa Lee for critical comments on the manuscript and the ABRC for providing cDNA. This work was supported by the National Science Foundation (PGRP BIO-2112056 to LCS), the National Institutes of Health (R35 GM136338 to LCS), and Duke Beyond the Horizons (to LCS) funds. This work was also supported by the Chan Zuckerberg Initiative DAF (an advised fund of Silicon Valley Community Foundation; grant 2023-332391; to JAJ), the National Institute of General Medical Sciences of the National Institutes of Health under Award Number R35GM155259 (to JAJ), and the National Science Foundation (NSF) through the Princeton University (PCCM) Materials Research Science and Engineering Center DMR-2011750 (to JAJ).

## Author Contributions

S.P., A.C. J.A.J. and L.C.S. designed the study. S.P. and C.B. performed microscopy and *in vivo* condensation assays. A.C. performed simulations and analyzed data. A.C. and S.P wrote the manuscript. L.C.S. and J.A.J. supervised the project.

## Competing Interests

L.C.S. is on the scientific advisory board of Prose Foods. All other authors declare no competing interests.

Supplementary Information is available for this paper.

## Materials and Correspondence

Correspondence and material requests should be addressed to Jerelle A. Joseph or to Lucia C. Strader.

## Supplementary Information

**Figure S1.**
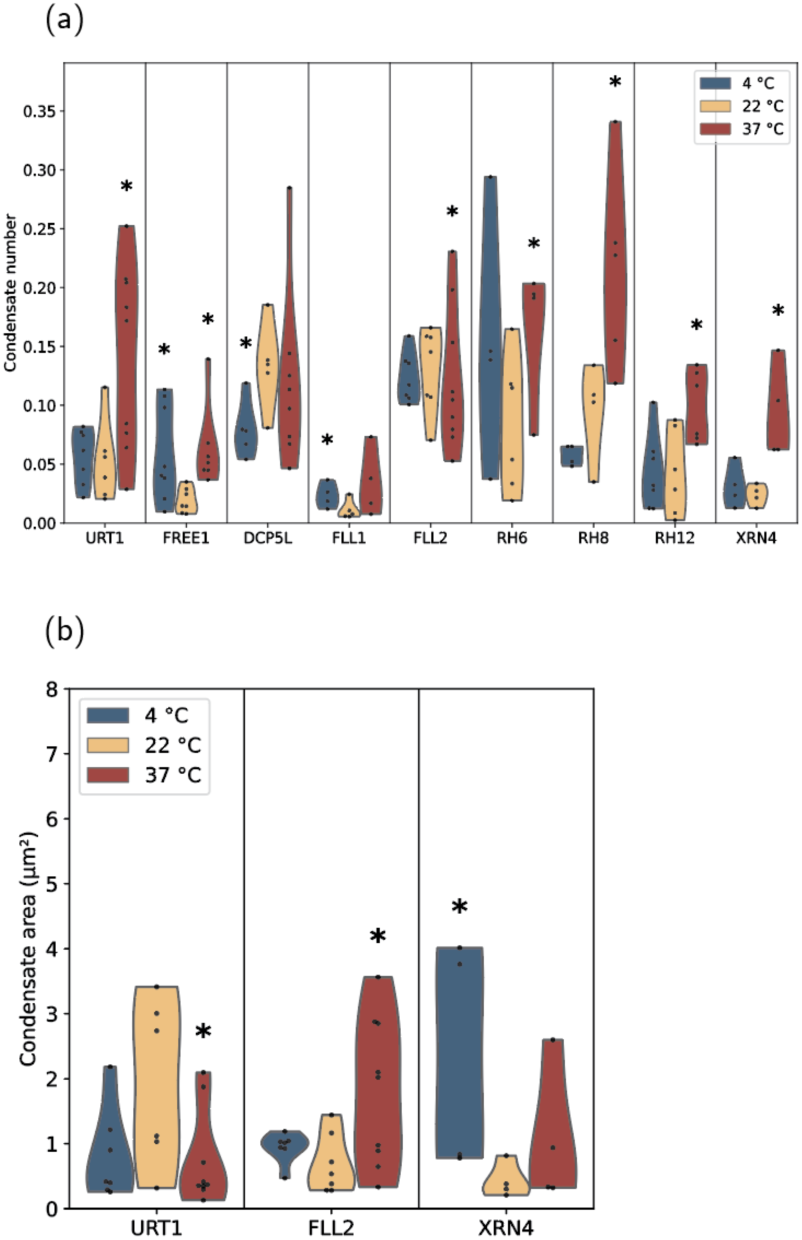
Cluster 1 protein from Figure 2 shows thermoresponsive change in condensate morphology. (a) Violin plot representing a significant change in condensate number (No. of condensate per area of protoplast) in each protoplast expressing cluster 1 proteins at 37°C, 22°C, and 4°C. (b) Violin plot representing a significant change in average area of condensates (µm^2^) in each protoplast expressing cluster 1 proteins at 37°C, 22°C, and 4°C.Each jitter represents data points from an individual protoplast.

**Figure S2.**
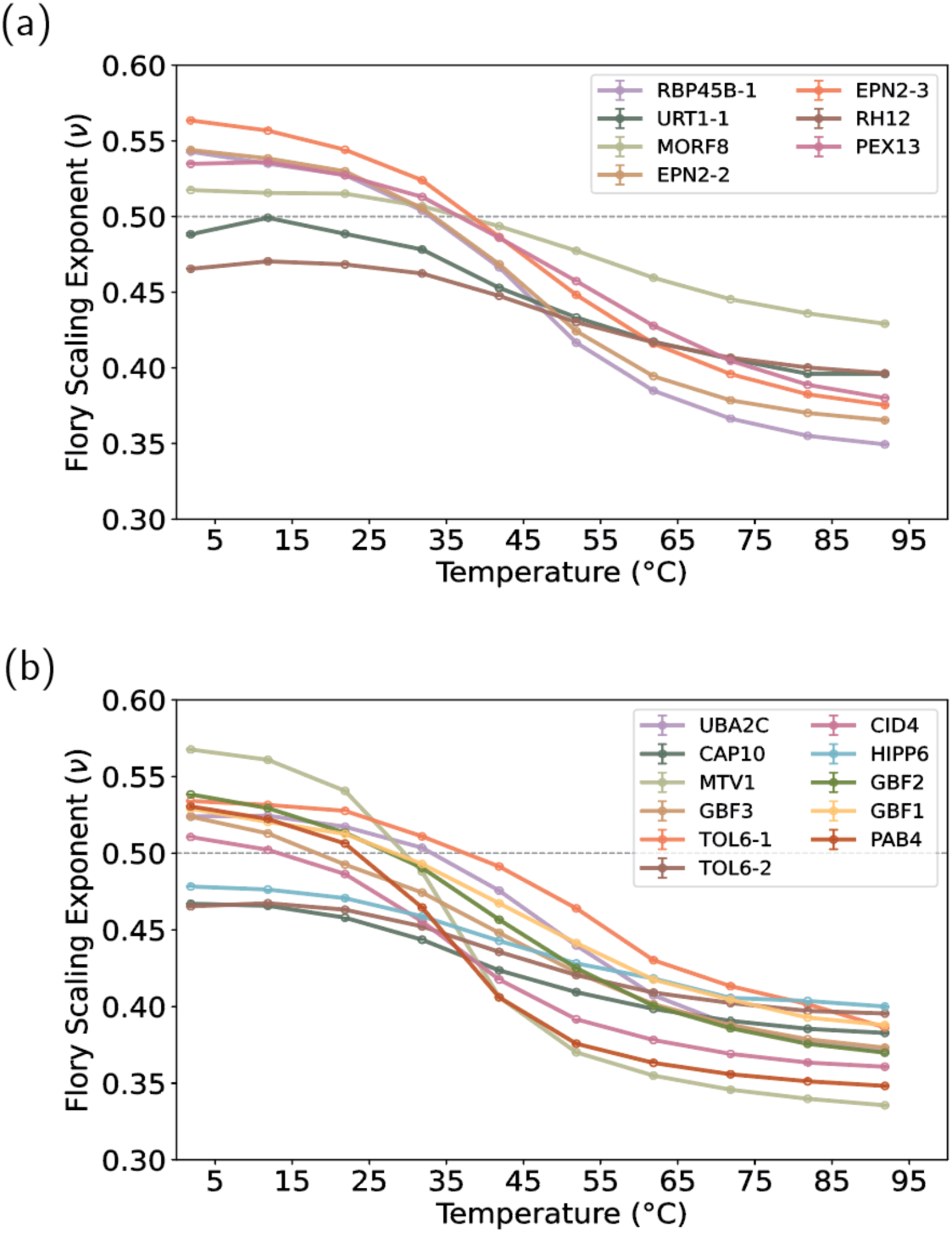
Single-chain simulation results for PLCD sequences. Single-chain simulations for PLCD sequences from Fig. 4c that are. (a) not sufficient or necessary for full-length protein phase separation and (b) sufficient for full-length protein phase separation. The dashed gray line marks the coil-to-globule transition. For sequences that exhibit a sharp increase in compactness below ν = 0.5, the transition temperature was defined as the point of steepest slope. Data sets were divided into three blocks, and the block means were used as the reported data points; error bars indicate the standard deviation across blocks. PLCD sequences from Fig. 3 that do not appear here are omitted because they did not display a discernible coil-to-globule trend under the conditions tested. For these sequences, ν remained approximately constant or increased across the simulated temperature range, preventing reliable identification of or indicating lack of an LCST transition.

**Figure S3.**
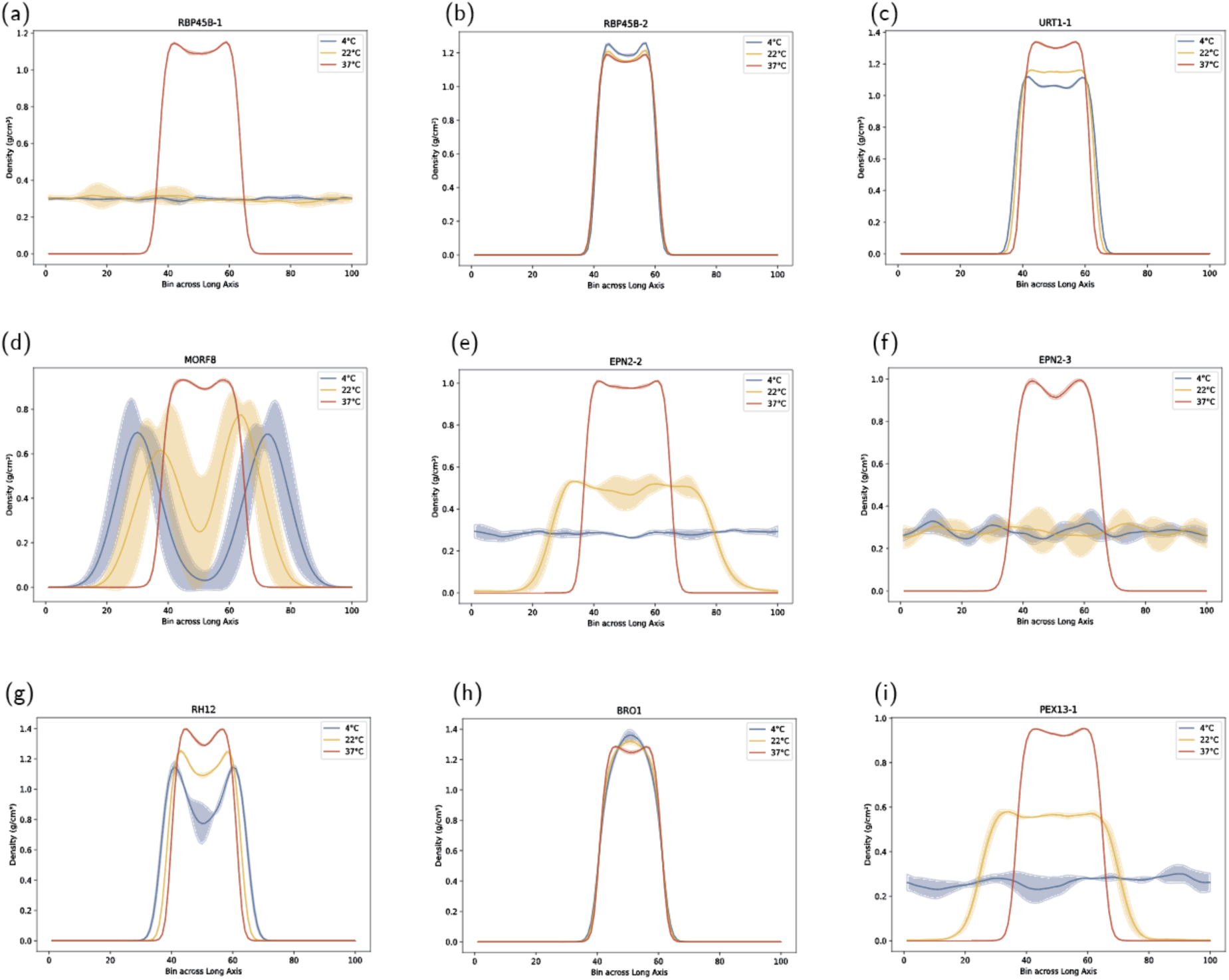
Density profiles to obtain dense-phase densities for PLCD sequences that are not necessary or sufficient for full-length protein phase separation. Direct coexistence simulations were performed at 4 °C, 22 °C, and 37 °C for all PLCD sequences analyzed in Fig. 3 to determine the dense-phase densities reported in Fig. 4d. To evaluate uncertainty, each trajectory was divided into three blocks; the block means define the data points, and the error bands represent the standard deviation.

**Figure S4.**
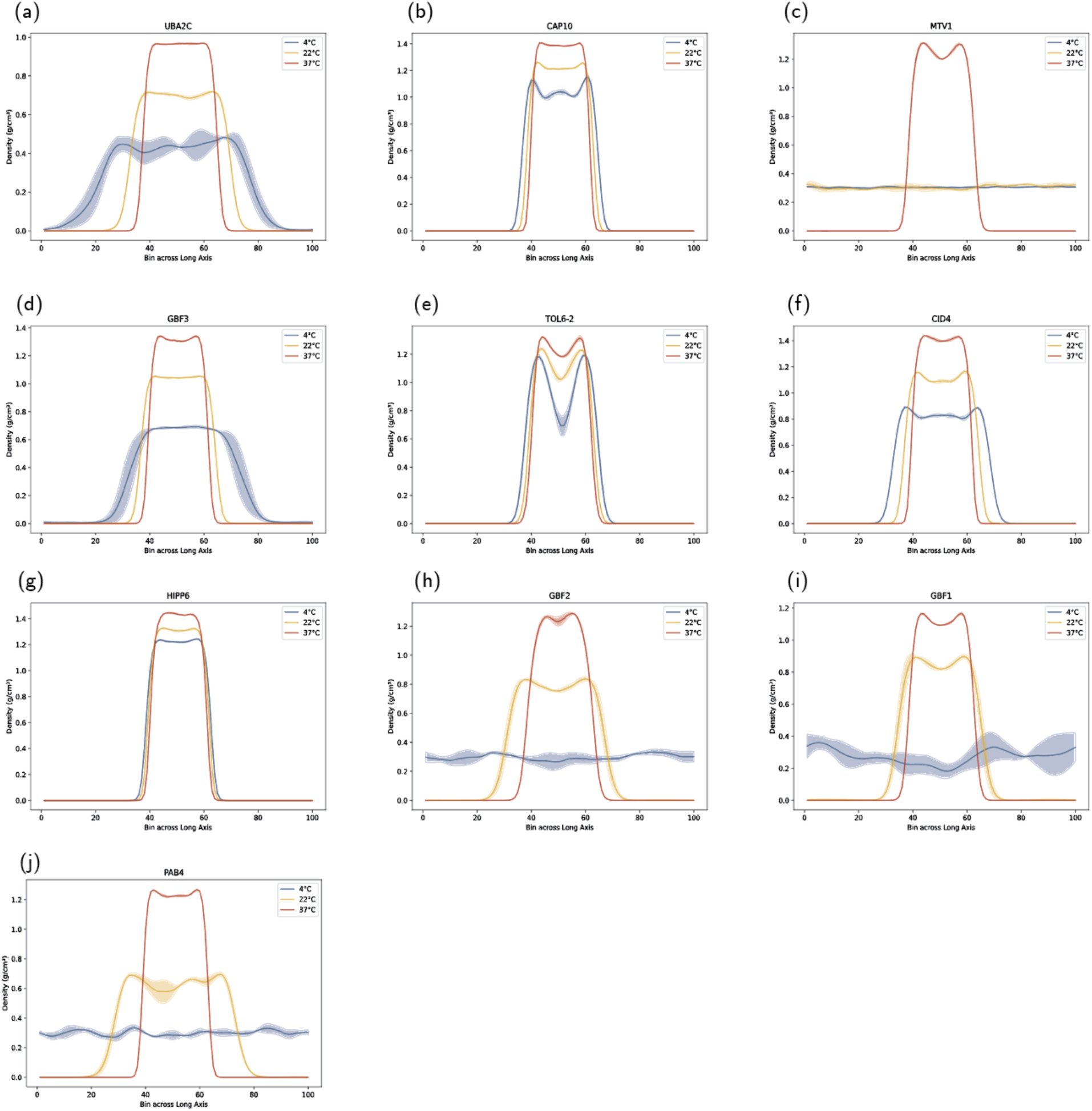
Density profiles to obtain dense-phase densities for PLCD sequences that are sufficient for full-length protein phase separation. Direct coexistence simulations were performed at 4 °C, 22 °C, and 37 °C for all PLCD sequences analyzed in Fig. 3 to determine the dense-phase densities reported in Fig. 4d. To evaluate uncertainty, each trajectory was divided into three blocks; the block means define the data points, and the error bands represent the standard deviation.

**Figure S5.**
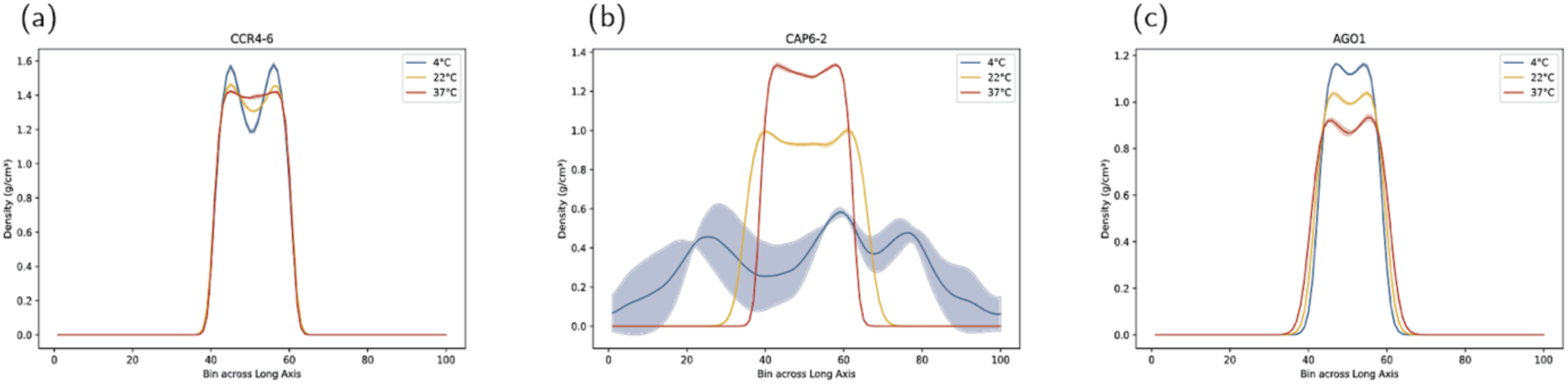
Density profiles to obtain dense-phase densities for PLCD sequences that are necessary for full-length protein phase separation. Direct coexistence simulations were performed at 4 °C, 22 °C, and 37 °C for all PLCD sequences analyzed in Fig. 3 to determine the dense-phase densities. To evaluate uncertainty, each trajectory was divided into three blocks; the block means define the data points, and the error bands represent the standard deviation.

**Figure S6.**
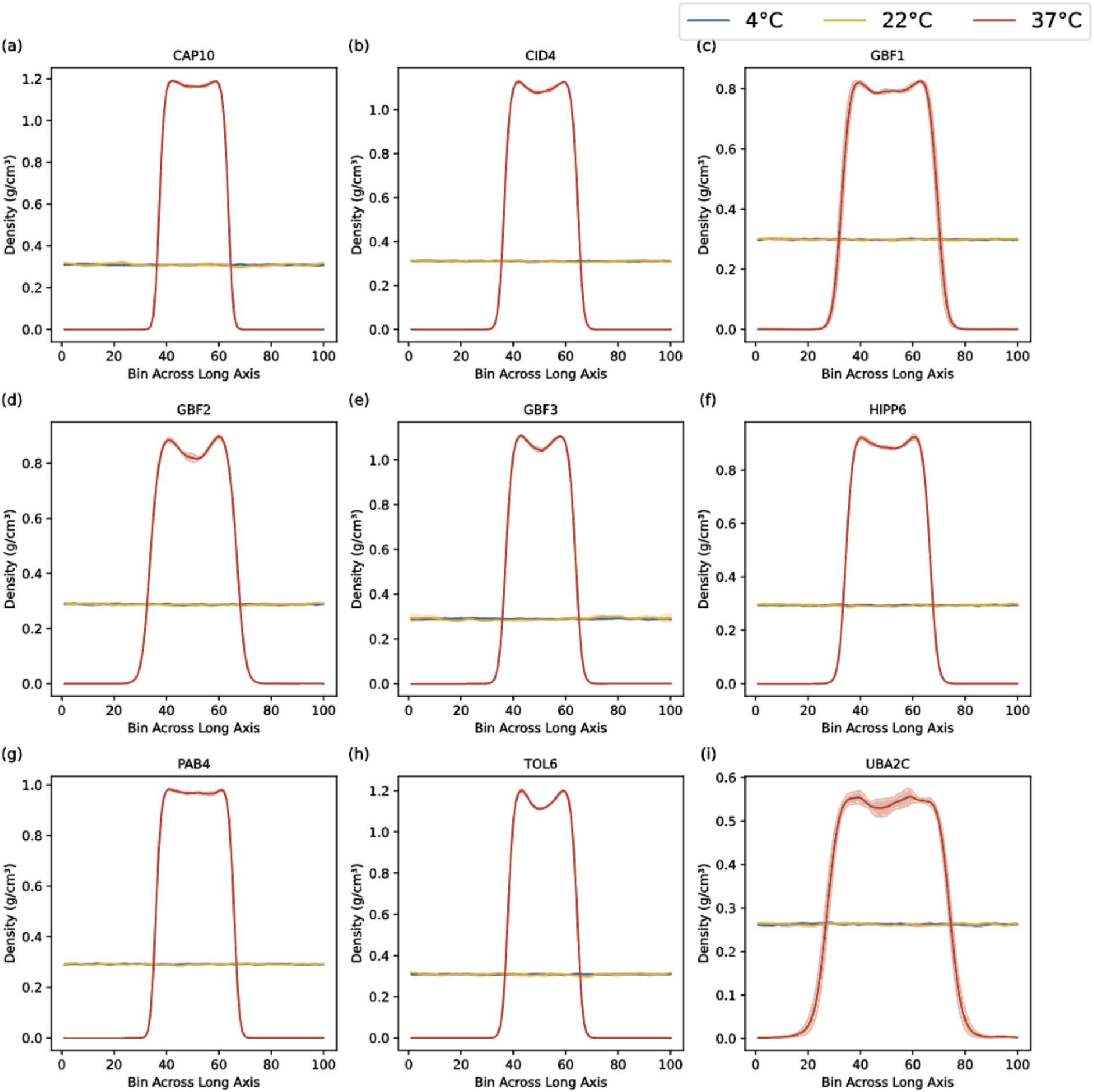
Density profiles to obtain dense-phase densities for proteins from Variant Set 1 (mutated polar or aromatic residues to aliphatic residues). Direct coexistence simulations were performed at 4 °C, 22 °C, and 37 °C for designed sequences in Fig. 5 to determine the dense-phase densities. To evaluate uncertainty, each trajectory was divided into three blocks; the block means define the data points, and the error bands represent the standard deviation.

**Figure S7.**
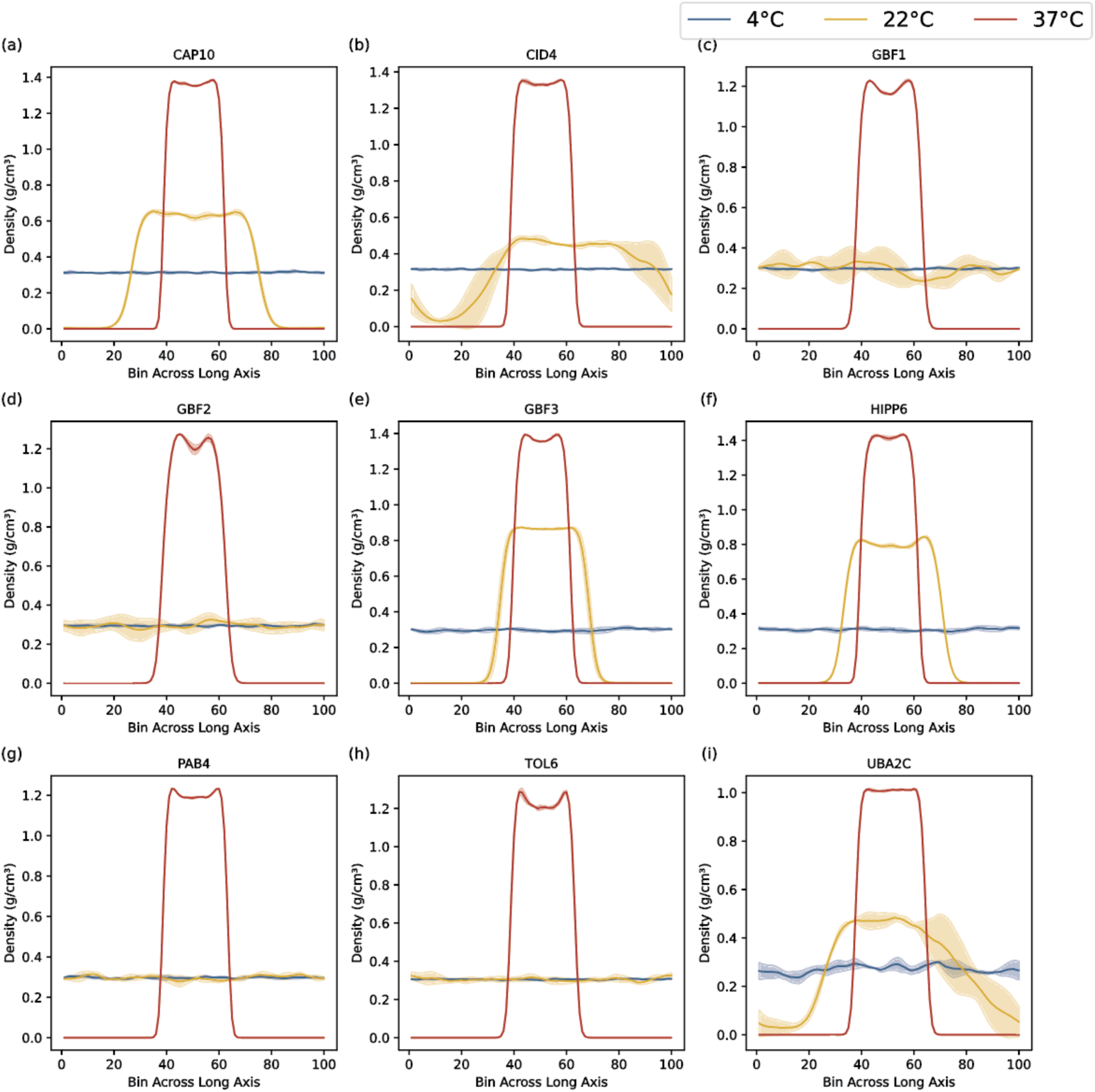
Density profiles to obtain dense-phase densities for proteins from Variant Set 2 (mutated polar residues to aliphatic residues). Direct coexistence simulations were performed at 4 °C, 22 °C, and 37 °C for designed sequences in Fig. 5 to determine the dense-phase densities. To evaluate uncertainty, each trajectory was divided into three blocks; the block means define the data points, and the error bands represent the standard deviation.

**Figure S8.**
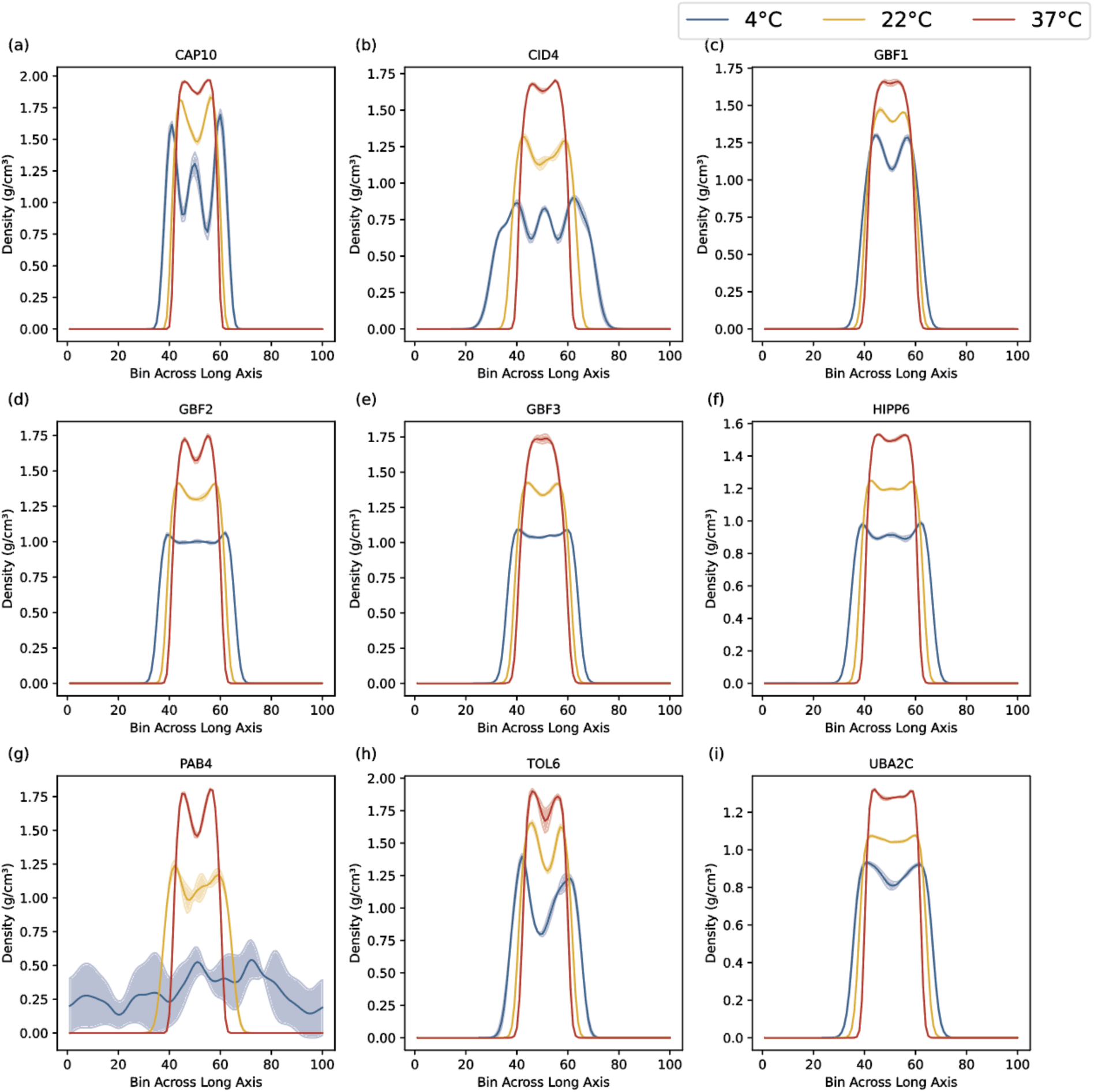
Density profiles to obtain dense-phase densities for proteins from Variant Set 3 (mutated polar residues to aliphatic or aromatic residues). Direct coexistence simulations were performed at 4 °C, 22 °C, and 37 °C for designed sequences in Fig. 5 to determine the dense-phase densities. To evaluate uncertainty, each trajectory was divided into three blocks; the block means define the data points, and the error bands represent the standard deviation.

## References

1. Casal, J.J. and S. Balasubramanian, Thermomorphogenesis. Annual Review of Plant Biology, 2019. 70(Volume 70, 2019): p. 321–346.

2. Hatfield, J.L. and J.H. Prueger, Temperature extremes: Effect on plant growth and development. Weather and Climate Extremes, 2015. 10: p. 4–10.

3. Zhao, C., et al., Temperature increase reduces global yields of major crops in four independent estimates. Proceedings of the National Academy of Sciences, 2017. 114(35): p. 9326–9331.

4. Xiong, J., et al., Emerging strategies to improve heat stress tolerance in crops. aBIOTECH, 2025. 6(1): p. 97–115.

5. Hayes, S., et al., Hot topic: Thermosensing in plants. Plant, Cell & Environment, 2020.

6. Ashraf, M., Thermotolerance in plants: Potential physio-biochemical and molecular markers for crop improvement. Environmental and Experimental Botany, 2021. 186: p. 104454.

7. Chung, B.Y.W., et al., An RNA thermoswitch regulates daytime growth in Arabidopsis. Nature Plants, 2020. 6(5): p. 522–532.

8. Jung, J.-H., et al., Phytochromes function as thermosensors in Arabidopsis. Science, 2016. 354(6314): p. 886–889.

9. Jung, J.-H., et al., A prion-like domain in ELF3 functions as a thermosensor in Arabidopsis. Nature, 2020. 585(7824): p. 256–260.

10. Haider, S., et al., Molecular mechanisms of plant tolerance to heat stress: current landscape and future perspectives. Plant Cell Rep, 2021. 40(12): p. 2247–2271.

11. Wang, Y.-X. and J.-H. Xu, The heat shock transcription factors regulate response mechanisms to abiotic stresses in plants. Crop Design, 2025. 4(3): p. 100109.

12. Xu, F., et al., Phase separation of GRP7 that is facilitated by FERONIA-mediated phosphorylation inhibits mRNA translation to modulate plant temperature resilience. Molecular Plant, 2024.

13. Peng, M., et al., Activation and memory of the heatshock response is mediated by Prion-like domains of sensory HSFs in Arabidopsis. Mol Plant, 2025.

14. Bohn, L., et al., The temperature sensor TWA1 is required for thermotolerance in Arabidopsis. Nature, 2024. 629(8014): p. 1126–1132.

15. Peng, J., Y. Yu, and X. Fang, Stress sensing and response through biomolecular condensates in plants. Plant Commun, 2025. 6(2): p. 101225.

16. Geng, P., et al., A thermosensor FUST1 primes heat-induced stress granule formation via biomolecular condensation in Arabidopsis. Cell Research, 2025. 35(7): p. 483–496.

17. Chen, D., et al., Integration of light and temperature sensing by liquid-liquid phase separation of phytochrome B. Molecular Cell, 2022. 82(16): p. 3015–3029.e6.

18. Londoño Vélez, V., et al., Landscape of biomolecular condensates in heat stress responses. Frontiers in Plant Science, 2022. 13.

19. Chantarachot, T. and J. Bailey-Serres, Polysomes, stress granules, and processing bodies: a dynamic triumvirate controlling cytoplasmic mRNA fate and function. Plant physiology, 2018. 176(1): p. 254–269.

20. Wu, J., et al., Solid-like condensation of MORF8 inhibits RNA editing under heat stress in Arabidopsis. Nature Communications, 2025. 16(1): p. 2789.

21. Wallace, E.W., et al., Reversible, Specific, Active Aggregates of Endogenous Proteins Assemble upon Heat Stress. Cell, 2015. 162(6): p. 1286–98.

22. Gu, J., et al., Advances in the structures, mechanisms and targeting of molecular chaperones. Signal Transduction and Targeted Therapy, 2025. 10(1): p. 84.

23. Legoux, M., J.P. Reichheld, and R. Merret, The role of post-translational modifications in the dynamics of cytoplasmic biomolecular condensates in plants. New Phytol, 2025. 248(4): p. 1692–1699.

24. Sabate, R., et al., What makes a protein sequence a prion? PLoS computational biology, 2015. 11(1): p. e1004013.

25. Li, X. and T. Kahveci, A Novel algorithm for identifying low-complexity regions in a protein sequence. Bioinformatics, 2006. 22(24): p. 2980–2987.

26. Molliex, A., et al., Phase separation by low complexity domains promotes stress granule assembly and drives pathological fibrillization. Cell, 2015. 163(1): p. 123–133.

27. Maristany, M.J., et al., Decoding phase separation of prion-like domains through data-driven scaling laws. eLife, 2025. 13: p. RP99068.

28. Riback, J.A., et al., Stress-Triggered Phase Separation Is an Adaptive, Evolutionarily Tuned Response. Cell, 2017. 168(6): p. 1028–1040.e19.

29. Chakrabortee, S., et al., Luminidependens (LD) is an Arabidopsis protein with prion behavior. Proceedings of the National Academy of Sciences, 2016. 113(21): p. 6065–6070.

30. Powers, S.K., et al., Nucleo-cytoplasmic partitioning of arf proteins controls auxin responses in Arabidopsis thaliana. Molecular cell, 2019. 76(1): p. 177–190. e5.

31. Lancaster, A.K., et al., PLAAC: a web and command-line application to identify proteins with prion-like amino acid composition. Bioinformatics, 2014. 30(17): p. 2501–2.

32. March, Z.M., O.D. King, and J. Shorter, Prion-like domains as epigenetic regulators, scaffolds for subcellular organization, and drivers of neurodegenerative disease. Brain Res, 2016. 1647: p. 9–18.

33. Gotor, N.L., et al., RNA-binding and prion domains: the Yin and Yang of phase separation. Nucleic Acids Research, 2020. 48(17): p. 9491–9504.

34. Garai, S., et al., Complex networks of prion-like proteins reveal cross talk between stress and memory pathways in plants. Frontiers in plant science, 2021. 12: p. 707286.

35. Alberti, S., et al., A Systematic Survey Identifies Prions and Illuminates Sequence Features of Prionogenic Proteins. Cell, 2009. 137(1): p. 146–158.

36. Maristany, M.J., et al., Decoding Phase Separation of Prion-Like Domains through Data-Driven Scaling Laws. 2025, eLife Sciences Publications, Ltd.

37. Blagojevic, A., et al., Heat stress promotes Arabidopsis AGO1 phase separation and association with stress granule components. Iscience, 2024.

38. Weber, C., L. Nover, and M. Fauth, Plant stress granules and mRNA processing bodies are distinct from heat stress granules. The Plant Journal, 2008. 56(4): p. 517–530.

39. Sheth, U. and R. Parker, Decapping and decay of messenger RNA occur in cytoplasmic processing bodies. Science, 2003. 300(5620): p. 805–8.

40. Chicois, C., et al., The UPF1 interactome reveals interaction networks between RNA degradation and translation repression factors in Arabidopsis. The Plant Journal, 2018. 96(1): p. 119–132.

41. Liu, S., C. Wang, and B. Zhang, Toward Predictive Coarse-Grained Simulations of Biomolecular Condensates. Biochemistry, 2025. 64(8): p. 1750–1761.

42. Chakravarti, A. and J.A. Joseph, Accurate prediction of thermoresponsive phase behavior of disordered proteins. Protein Science, 2025. 34(10): p. e70284.

43. Martin, E.W. and T. Mittag, Relationship of Sequence and Phase Separation in Protein Low-Complexity Regions. Biochemistry, 2018. 57(17): p. 2478–2487.

44. Quiroz, F.G. and A. Chilkoti, Sequence heuristics to encode phase behaviour in intrinsically disordered protein polymers. Nature Materials, 2015. 14(11): p. 1164–1171.

45. Wu, F.-H., et al., Tape-Arabidopsis Sandwich-a simpler Arabidopsis protoplast isolation method. Plant methods, 2009. 5(1): p. 1–10.

46. Kostylev, M., et al., Cloning Should Be Simple: Escherichia coli DH5α-Mediated Assembly of Multiple DNA Fragments with Short End Homologies. PLoS One, 2015. 10(9): p. e0137466.

47. Allen, J.R., et al., An improved toolkit of gateway-and gibson assembly-compatible vectors for protoplast transfection and agrobacterium-mediated plant transformation. BMC Research Notes, 2025. 18(1): p. 149.

48. Pedregosa, F., et al., Scikit-learn: Machine Learning in Python. J. Mach. Learn. Res., 2011. 12(null): p. 2825–2830.

